# CREB-family transcription factors and vasopressin-mediated regulation of *Aqp2* gene transcription

**DOI:** 10.1101/2025.03.10.642395

**Authors:** Adrian Rafael Murillo-de-Ozores, Lihe Chen, Shuo-Ming Ou, Euijung Park, Shaza Khan, Viswanathan Raghuram, Chin-Rang Yang, Chung-Lin Chou, Mark A. Knepper

**Affiliations:** Epithelial Systems Biology Laboratory, Systems Biology Center, National Heart, Lung, and Blood Institute, National Institutes of Health, Bethesda, MD

**Keywords:** vasopressin, water balance, CRISPR-Cas9, CREB, ATF1, CREM

## Abstract

**Background:** Water homeostasis is regulated by the peptide hormone arginine vasopressin (AVP), which promotes water reabsorption in the renal collecting duct. The regulation of *Aqp2* gene transcription is a key mechanism through which AVP modulates water transport as disruption of this mechanism leads to water balance disorders. Therefore, an important goal is to understand the regulatory processes that control *Aqp2* gene transcription. While CREB (CREB1) has been proposed as the primary transcription factor responsible for *Aqp2* transcription, recent evidence challenges this view, suggesting that other CREB-like transcription factors, including ATF1 and CREM, may play a role.

**Methods:** We employed the CRISPR/Cas9 gene-editing system to delete *Atf1*, *Creb1*, and *Crem* in mpkCCD cells, an immortalized mouse collecting duct cell line. These cell lines were then exposed to the vasopressin analog, dDAVP, to assess the role of these transcription factors in regulating *Aqp2* expression. AQP2 protein levels were measured by immunoblotting and RNA-seq was used to analyze changes in *Aqp2* mRNA abundance, as well as other transcriptomic changes.

**Results:** Deletion of all three transcription factors (ATF1, CREB1, and CREM) led to a significant reduction in the vasopressin-induced upregulation of AQP2 protein, confirming their role in regulating *Aqp2* expression. RNA-seq data showed that *Aqp2* mRNA levels mirrored changes in protein abundance, supporting the idea that these transcription factors affect *Aqp2* transcription. Rescue experiments in triple knockout cells showed that expressing any of the three transcription factors restored the response to vasopressin.

**Conclusions:** Our findings demonstrate that ATF1, CREB1, and CREM have redundant roles in regulating *Aqp2* transcription. Based on these results and prior data, we propose that these CREB-family transcription factors may regulate *Aqp2* gene transcription indirectly by controlling the expression of additional unidentified transcription factors.

**Key Points:**

- **CREB-family transcription factors (ATF1, CREB1, and CREM) were deleted in mpkCCD cells to assess their roles in *Aqp2* gene transcription**
- **CRISPR/Cas9 knockout of all three transcription factors strongly reduced the ability of vasopressin to increase AQP2 mRNA and protein**
- **Re-expression of any of the three restored the vasopressin response indicating redundant roles of the three transcription factors**

## Introduction

Water homeostasis is regulated by the peptide hormone arginine vasopressin (AVP). This molecule is produced in the hypothalamus and released by the neurohypophysis in response to increased blood osmolality. AVP binding to the V2 receptor (V2R) in the principal cells of the collecting duct regulates sodium and water transport proteins^1-7^. Alterations in the function of AVP or its downstream signaling result in dysregulation of water balance, characterized either by excess water excretion resulting in dilutional hyponatremia (e.g. diabetes insipidus) or by excess water retention (e.g. syndrome of inappropriate antidiuresis (SIAD)^8-12^.

In the collecting duct, AVP promotes the upregulation of aquaporin 2 (AQP2)-mediated water reabsorption by two mechanisms: 1) a short-term effect on trafficking of AQP2-containing membrane vesicles to and from the apical plasma membrane^13^ and 2) long-term regulation by increasing AQP2 protein abundance^14^. The second mechanism is largely attributed to increased *Aqp2* gene transcription in response to V2R signaling^15,16^. Studies in animal models of water balance disorders (both diabetes insipidus syndromes and dilutional hyponatremia) have demonstrated that the dysregulation is due to failure of the second mechanism, i.e. loss of regulation of *Aqp2* gene transcription^8^. Therefore, an important goal is to understand the regulatory processes that control *Aqp2* gene transcription. V2R is a G_α_s-coupled receptor that promotes an increase in intracellular cAMP levels and therefore, protein kinase A (PKA) activation. Deletion of both catalytic subunits of PKA (PKA double KO) led to an almost complete loss of vasopressin-dependent *Aqp2* gene expression^17^.

Several review papers have proposed that PKA regulates *Aqp2* gene transcription by phosphorylating the transcription factor CREB (also known as <u>C</u>yclic AMP-<u>R</u>esponsive <u>E</u>lement-<u>B</u>inding Protein <u>1</u>: CREB1)^5,8,18,19^. Deletion of the cAMP response element (CRE) in constructs containing the *Aqp2* promoter has been shown to decrease promoter activity in response to cAMP analogues in non-collecting duct cells^15,20,21^. Additional experiments support the role of CREB in regulation of *Aqp2* gene transcription, including immunoblotting showing that vasopressin promotes CREB phosphorylation, as well as electrophoretic mobility shift assays suggesting that CREB might bind to a CRE in the *Aqp2* promoter *in vitro*^21^. However, ChIP-seq studies in a collecting duct cell line that endogenously expresses AQP2 (mpkCCD cells) have not found CREB binding sites within 390 kb of the transcription start site of the *Aqp2* gene despite showing strong binding to the promoters of several previously identified CREB target genes^22^, suggesting that CREB’s role in regulating *Aqp2*, if any, might be indirect.

CREB1 (Gene symbol: *Creb1*) is a member of a family of PKA-activated transcription factors, which also includes CREM (Gene symbol: *Crem*) and ATF1 (Gene symbol: *Atf1*)^23^. All three are plausible candidates to mediate vasopressin’s effect in increasing *Aqp2* gene transcription. They are basic leucine zipper (bZIP) transcription factors characterized by a “kinase-inducible domain” (KID) that contains a conserved phosphorylation site that can be a target of PKA or other basophilic kinases^24^. Phosphorylation of this site enhances CREB, ATF1 or CREM activity by promoting their interaction with transcriptional co-activators such as CREBBP and/or EP300^23,25^. CREM is expressed as either the full length form and a short isoform called “ICER” which lacks the KID domain^26^. Previously, we have described a Bayesian analysis to rank candidate transcription factors that might bind near the *Aqp2* gene promoter or a putative enhancer and regulate its transcription^27^. This unbiased approach identified ATF1 as one of the top candidates.

In this study, we employed the CRISPR/Cas9 system to delete *Atf1*, *Creb1*, and *Crem* in order to investigate if they play causal roles in the upregulation of AQP2 expression in response to the vasopressin analog, dDAVP. By analyzing the effects of these deletions, we aim to clarify the contribution of these transcription factors (if any) in *Aqp2* gene expression and to identify ATF1/CREB1/CREM transcriptional targets in mpkCCD cells.

## METHODS

### Cell lines

Immortalized mpkCCD clone11-38^28^ was transfected with pCMV-Cas9 with either GFP or RFP plasmids (Sigma) with sgRNAs directed against *Atf1*, *Creb1* or *Crem* genes, using Lipofectamine 3000 (Invitrogen) following the manufacturer’s instructions. sgRNAs sequences were as follows: *Atf1:* 5’-GATACTCGTCCCGAGCAACCAGG-3’, 5’-TCCAAGCACGGATGGAGTGCAGG-3’, *Creb1* 5’-CTGGCTAACAATGGTACGGATGG-3’, 5’-CAATGGTACGGATGGGGTACAGG-3’ and *Crem* 5’-CAGTAGTAGGAGCTCGGATCTGG-3’, 5’-GGAGCTCGGATCTGGTAAGTTGG-3’. Non-targeting sgRNAs were used as controls: 5’-CGCGATAGCGCGAATATATT-3’, 5’-GCGCGATAGCGCGAATATAC-3’. GFP or RFP positive cells were subjected to single-cell sorting into 96-well plates using a BD FACSAria II cell sorter. Cells were progressively expanded and DNA and protein was obtained. ATF1, CREB1 or CREM protein expression was evaluated by immunoblotting. Cell clones with absence of the target protein were further confirmed as knockouts by Sanger sequence to detect genomic indel mutations in the target genes. PKA-double KO mpkCCD cells have been described previously^17^.

### Cell culture

Cells were maintained in DMEM/F-12 media with 2% Fetal Bovine Serum and supplemented with 5 μg/mL insulin, 50 nM dexamethasone, 1 nM triiodothyronine, 10 ng/mL epidermal growth factor, 60 nM sodium selenite, 5 μg/mL transferrin; all from Sigma). In order to polarize cells, they were grown in permeable membrane supports (Transwell, Corning #3450) and grown with complete media for 4 days, prior to changing to simple media (DMEM/F-12 media only supplemented with 50 nM dexamethasone, 60 nM sodium selenite and 5 μg/mL transferrin) for 3 days, either with or without 0.1 nM dDAVP (1-desamino-8-D-arginine-vasopressin).

### Generation of anti-ATF1 antibody

A peptide corresponding to amino acids 73 to 90 of mouse ATF1 was selected using AbDesigner^29^, synthesized with an N-terminal Cys (NH_2_-CSEDTRGRKGEGENPSISA-COOH), HPLC-purified and KLH-conjugated. Rabbits were immunized with this peptide using a standard protocol. Antibody was affinity-purified with a peptide-coupled column (Pierce, SulfoLink Kit). The antibody’s specificity was confirmed by Western Blot against the peptide, as well as comparison between cell lysates from control and *Atf1*-KO cells.

### Western Blotting

Cells were washed with ice-cold Dulbecco’s Phosphate-Buffered Saline (DPBS) and then lysed with lysis buffer (1.5% SDS, 10mM Tris pH 6.8) containing protease and phosphatase inhibitor (78441, Thermo Fisher Scientific). Protein concentration was quantified by BCA assay (Pierce). 10-25 μg of protein were used for SDS-PAGE using 12% Criterion TGX gels (5671045, Bio-Rad). Proteins were transferred to nitrocellulose membranes, which were blocked (Intercept® (PBS) Blocking Buffer) for 1 hour at room temperature. Primary antibodies were incubated overnight at 4 °C. After washing, secondary antibodies were incubated for 1 hour at room temperature. After washing again, images were acquired by Li-COR Odyssey System. Band intensities were analyzed with ImageJ. The antibodies used were: anti-AQP2 (Knepper’s lab, K5007^30^), anti-phospho-CREB-S133 (Cell Signaling, 9198), anti-ATF1 (this study), anti-CREB (Sigma, 04-767), anti-FLAG (Cell Signaling, 14793), anti-rabbit (Li-COR, IRDye® 680RD).

### RNA Isolation and Sequencing

Total RNA was isolated from three biological replicates of each corresponding group using a Direct-zol RNA MiniPrep Plus Kit (Zymo Research) following the manufacturer’s protocol. First strand cDNA was prepared using 40 ng of total RNA using SMART-Seq® mRNA Kit, followed by cDNA amplification. After purification of amplified cDNA using the AMPure XP beads (A63880, Beckman Coulter), the concentration of the synthesized cDNA was examined by Qubit dsDNA HS DNA assay kit (Q32851, Invitrogen). One nanogram of the synthesized cDNAs were fragmented and tagged with index primers (FC-131-1024, Nextera XT DNA library Preparation Kit, Illumina) following the manufacturer’s protocol. RNA-seq was performed by 50-bp paired-end NovaSeq (Illumina). Ten samples were loaded in each lane. Raw sequencing reads were aligned by STAR 2.7.10 b to the mouse reference genome (Ensembl genome 106). Transcript per million and expected counts were generated by RSEM 1.3.1. Expected counts were used for differential gene expression analysis using edgeR 3.38.4.

### Lentiviral transduction

Mouse *Atf1* (MR223254), *Creb1* (MR204788) and *Crem* (MR224371) ORF clones (OriGene Technologies) were used for PCR amplification and insertion into pWPXLd plasmid using In-Fusion cloning (Takara Bio) following manufacturer’s instructions. This construct is designed for producing FLAG-tagged proteins, as well as GFP with a self-cleaving peptide P2A in the middle. Lentix293T cells were transfected (Lipofectamine 3000, Thermo Fisher Scientific) with the corresponding pWPXLd plasmid as well as psPAX2 and pMD2.G plasmids (Addgene 12260 and 12259). Lentivirus-containing media was collected at 48 and 72 hours post-transfection, pooled together, centrifuged at 500g for 5 min and filtered (0.45 μm) to eliminate cellular debris. Aliquots were stored at -80 °C.

For viral transduction of mpkCCD cells, 50,000 cells were seeded along with the corresponding lentivirus (MOI ∼0.2) in the presence of 8 μg/mL polybrene (Millipore TR-1003-G). After 48 hours, GFP-positive cells were sorted using a BD FACSAria II cell sorter, expanded and used for polarization experiments.

### Bioinformatics and statistics

Most analyses were carried out using Microsoft Excel (https://www.microsoft.com/en-us/microsoft-365/excel) and R software (https://www.r-project.org/). The Database for Annotation, Visualization and Integrated Discovery (DAVID) (https://david.ncifcrf.gov/, RRID:SCR_001881) was used to identify gene set enrichment on the gene ontology database. The enrichment of specific gene ontology biological processes was defined by statistical evaluation using Fisher Exact test (p value < 0.05).

## RESULTS

RNA-seq data (**Figure 1A, 1B and 1C**) show that all three CREB-family transcription factors are expressed both in native mouse CCDs^31^ and mpkCCD (new data). Prior single-cell RNA-seq studies have demonstrated the presence of all three transcription factors in collecting duct principal cells as well as alpha and beta intercalated cells^32^. In a prior study, we produced mpkCCD-derived cell lines in which both catalytic genes (PKA catalytic-α and PKA catalytic-β) were deleted by CRISPR-Cas9^17^. We used these cells to characterize the response to dDAVP exposure (**Figure 1D, 1E, 1F**). As seen before^33^, a large increase in AQP2 protein abundance was seen after long-term exposure to dDAVP (**Figure 1D**). A phospho-specific antibody that recognizes the KID domain in CREB1 and ATF1 demonstrated a transient increase in phosphorylation of both transcription factors, which preceded the increase in AQP2 abundance (**Figure 1E and 1F**). The increases in AQP2 protein abundance, phosphorylated CREB1 and phosphorylated ATF1 were all ablated in the PKA double KO cells. Interestingly, the baseline phosphorylation of CREB1 and ATF1 was not eliminated in the PKA double KO mpkCCD cell lines indicating that some other basophilic protein kinase can also phosphorylate the KID domain, consistent with prior findings^34,35^. Additional immunoblotting of synthetic peptides corresponding to the non-phosphorylated and phosphorylated forms of the KID domain of CREB1 (**Figure 1G**) shows no recognition of the non-phosphorylated peptide by the phospho-specific antibody. Overall, the data in Figure 1 shows that mpkCCD cells provide a suitable model for investigation of the role of the three CREB-family transcription factors (if any) in regulation of AQP2 protein abundance.

**Figure 1.**
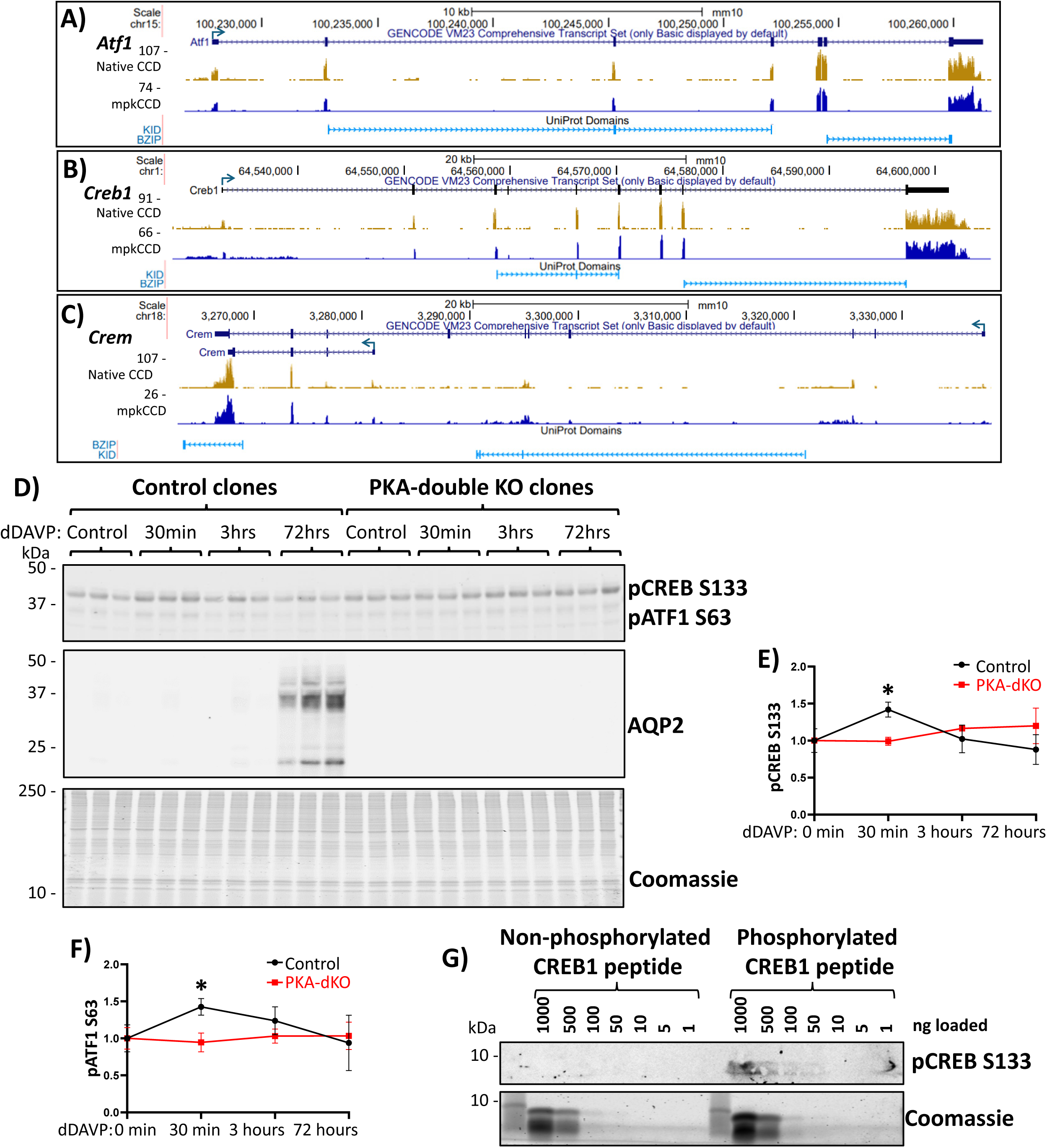
mpkCCD cell line as a model to study CREB-family transcription factors. RNA-seq data from native mouse collecting duct (CCD) and mpkCCD cells (this study) shows reads corresponding to the genes coding the three CREB-family transcription factors, (**A**) ATF1, (**B**) CREB, and (**C**) CREM. In panel C, two representative isoforms are shown for CREM, matching to full-length CREM (containing KID domain) and a shorter isoform lacking KID domain which corresponds to ICER. (**D**) Immunoblotting for pCREB, AQP2 and staining with Coomassie blue in successive sub-panels. Lysates were isolated from control and PKA-knockout mpkCCD cells treated with 0.1nM dDAVP for different times. First subpanel shows anti-pCREB antibody is able to detect both CREB (band at ∼40 kDa) and ATF1 (∼35 kDa). Second subpanel shows AQP2 abundance is considerably higher in control cells after 72 hours dDAVP incubation, but not in PKA-knockout cells. (**E and F**) Quantification of band intensities show that CREB and ATF1 phosphorylation is increased after 30 min dDAVP incubation in control cells but not in PKA-knockout cells. However, both CREB and ATF1 phosphorylation is still detectible in these cells. (**G**) Immunoblotting of CREB-peptide containing a serine or phospho-serine at position 133. Phospho-specific antibody used previously can only detect phosphorylated peptide.

### CRISPR-Cas9 deletion of ATF1

Our previous Bayesian analysis predicted a role for the KID domain transcription factor ATF1 in regulation of *Aqp2* gene transcription^27^. To test the hypothesis that ATF1 alone can mediate vasopressin’s effect to increase *Aqp2* gene transcription, we deleted ATF1 in mpkCCD cells using CRISPR-Cas9 (**Figure 2**). We made a new peptide-directed antibody for ATF1 protein (see ***Methods***). Immunoblots with this antibody revealed that control clones express ATF1 protein as a single band of ∼35 kD, while the ATF1-knockout (KO) cell lines do not show this band (**Figure 2A**). The control clones exhibited a strong increase in AQP2 protein abundance in response to dDAVP. However, the ATF1-KO clones also showed responses of similar magnitude. This implies that deletion of ATF1 by itself is insufficient to prevent vasopressin’s effect to increase *Aqp2* gene expression. Interestingly, the abundance of CREB1 was increased in the ATF1-KO cells (**Figure 2A**) both in vehicle-treated cells (**Figure 2B**) and in dDAVP-treated cells (**Figure 2C**). To address whether deletion of ATF1 alters the abundance of the *Aqp2* transcript or any other mRNA, we carried out RNA-seq in control cells and ATF1-KO cells, both in the presence of dDAVP (**Figure 2D**). A spreadsheet of all data values can be found at https://esbl.nhlbi.nih.gov/Databases/CREB-family/DataSets.html (Supplemental Spreadsheet 1). *Aqp2* mRNA was not decreased in the ATF1-KO cells (green point) and only a few transcripts showed significant changes. Transcripts that were decreased in the ATF1-KO cells included *Adh1* (alcohol dehydrogenase 1), which was found in previous studies to be strongly upregulated by vasopressin^16,36^. Otherwise, none of the responding transcripts were previously seen to be regulated by vasopressin. In particular, we did not see a change in CREB1 mRNA (yellow point), suggesting that the increase in CREB1 protein abundance in ATF1 KO cells is post-transcriptional.

**Figure 2.**
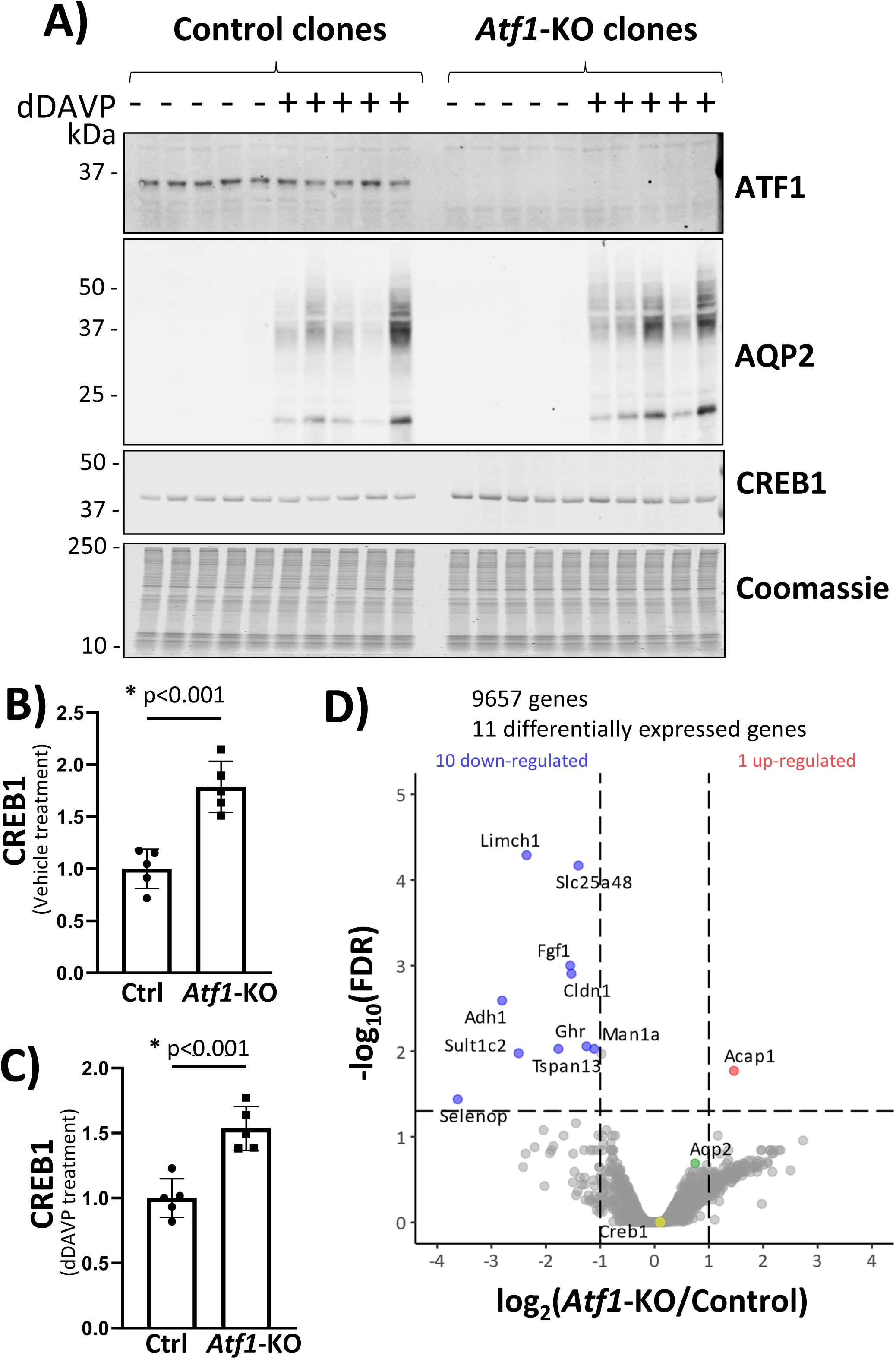
CRISPR-Cas9 deletion of ATF1 in mouse mpkCCD cells. (**A**) Immunoblotting for ATF1, AQP2, CREB1 and staining with Coomassie blue in successive sub-panels. Observations were made in cells exposed to dDAVP (dDAVP +) of vehicle (dDAVP -) for 72 hours after confluence. First subpanel shows loss of ATF1 protein in ATF1-KO lines compared with control clones. Second subpanel shows lack of detectible AQP2 in absence of dDAVP and a very large increase in AQP2 protein with dDAVP exposure. Third panel shows immunoblot for CREB1 showing a single band at ∼40 kDa. (B) Densitometry for CREB1 shows significant increase in CREB1 in ATF1-KO cells in absence of dDAVP. (C) Densitometry for CREB1 shows significant increase in CREB1 in ATF1-KO cells in the presence of dDAVP. (D) RNA-seq volcano plot showing transcripts increased or decreased according to threshold values: absolute value of log_2_(*Atf1*-KO/Control) greater than 1 and -log_10_(FDR-adjusted P value) greater than 1.301. All cells were grown in 0.1 nM dDAVP for 72 hours after confluence. Note that *Aqp2* (green point) or *Creb1* (yellow point) mRNAs were not significantly changed.

### ATF1/CREB1 double KO cells

Although ChIP-seq studies revealed a lack of CREB1 binding sites within 390 kb of the *Aqp2* gene transcriptional start site^22^, CREB1 could still regulate *Aqp2* gene transcription indirectly by controlling the abundance of some other transcriptional regulator. Accordingly, we have used CRISPR-Cas9 to create different ATF1/CREB1 double KO clones (see ***Methods***) (**Figure 3**). Figure 3A (first two subpanels) shows that ATF1 and CREB1 protein were both undetectable in the double KO clones. Immunoblotting for AQP2 confirmed that the ATF1 single KO cells respond to dDAVP with a large increase in AQP2 protein abundance (**Figure 3A**, third subpanel). The ATF1/CREB1 double KO cells were also found to respond to dDAVP with an increase in AQP2 protein that was approximately equal to that seen in the ATF1 single KO cells (**Figure 3A and 3B**). To address whether deletion of both ATF1 and CREB1 alters the abundance of the *Aqp2* transcript or any other mRNA, we carried out RNA-seq in control cells and ATF1/CREB1 double KO cells (**Figure 3C**). A spreadsheet of all data values can be found at https://esbl.nhlbi.nih.gov/Databases/CREB-family/DataSets.html (Supplemental Spreadsheet 2). Again there was no significant change in the abundance of *Aqp2* mRNA (green point). However, the overall response to the double deletion was much more substantial with more transcripts exhibiting significant changes, which on an average were larger than seen in the ATF1 single KO cells. Included in the list of increased transcripts were mRNAs that code for two regulators of mineralocorticoid responses in collecting duct principal cells, namely *Sgk1* and *Hsd11b2*. In addition, there was a significant increase in the mRNA that codes for the third KID domain transcription factor, CREM (purple point). Mapping the CREM reads on a genome browser (**Figure 3D**) shows increases in all exons indicating that the full length isoform is upregulated in the ATF1/CREB1 double KO cells.

**Figure 3.**
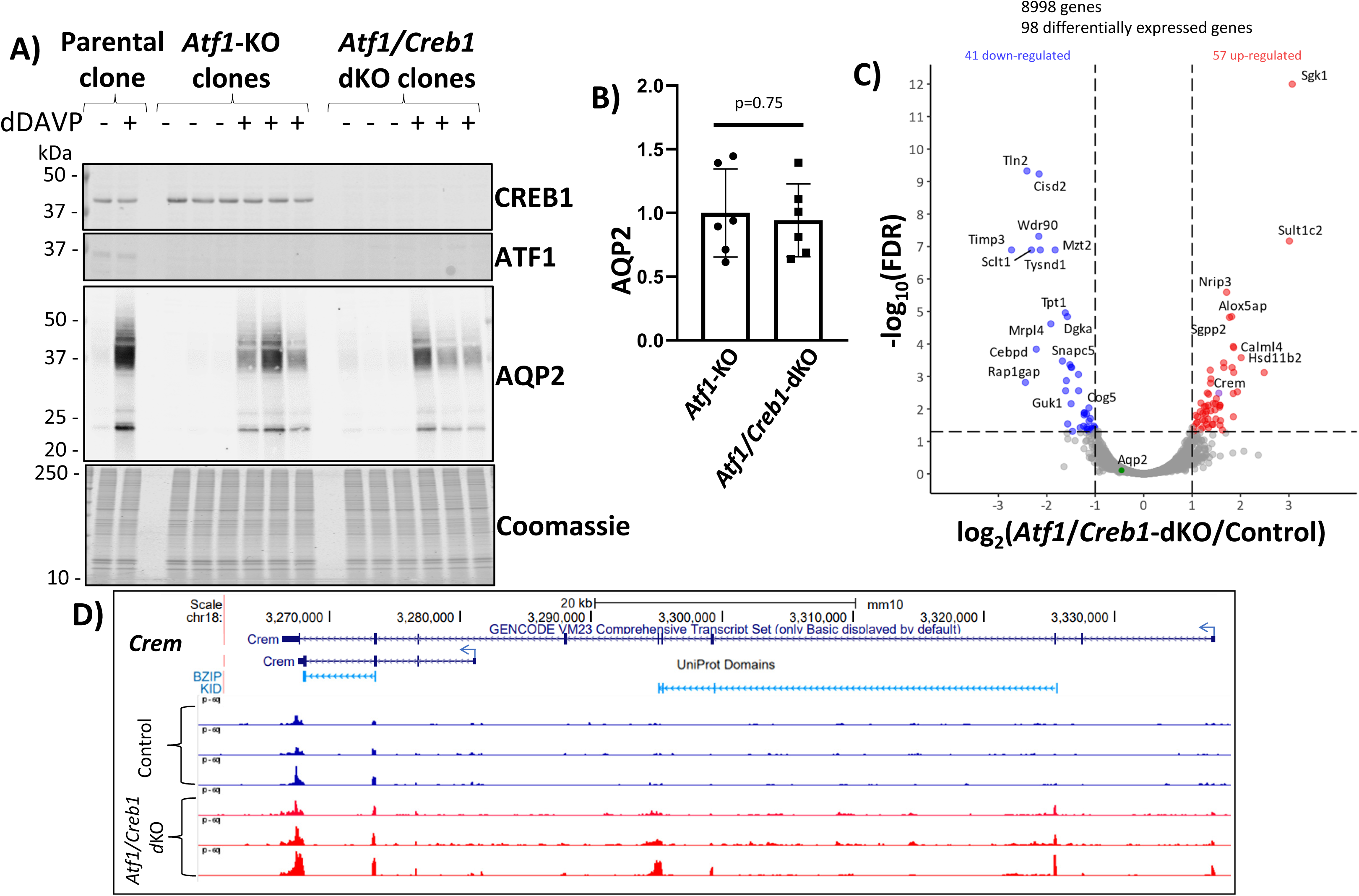
CRISPR-Cas9 deletion of ATF1 and CREB1 in mouse mpkCCD cells. (**A**) Immunoblotting for CREB1, ATF1 and AQP2 and staining with Coomassie blue in successive sub-panels. Observations were made in cells exposed to dDAVP (dDAVP +) of vehicle (dDAVP -) for 72 hours after confluence. The first two subpanels show that ATF1 is detectible only in the parental cells and CREB1 is only detectible in parental cells and ATF1 single KO cells. The third subpanel shows lack of detectible AQP2 in the absence of dDAVP in parental, ATF1-sKO and ATF1/CREB1 dKO cells and shows a very large increase in AQP2 protein with dDAVP exposure in all three genotypes. (**B**) Densitometry for AQP2 in dDAVP-exposed ATF1 sKO and ATF1/CREB1 dKO shows that deleting CREB1 on top of ATF1 does not substantially diminish the ability of dDAVP to increase AQP2 abundance. (**C**) RNA-seq volcano plot showing transcripts increased or decreased in ATF1/CREB1 dKO cells versus unmodified (control) cells according to threshold values: absolute value of log_2_(*Atf1/Creb1*-dKO/Control) greater than 1 and -log_10_(FDR-adjusted P value) greater than 1.301. All cells were grown in 0.1 nM dDAVP for 72 hours after confluence. Note that *Aqp2* (green point) mRNA was not significantly changed, while *Crem* (purple point) is upregulated. (**D**) RNA-seq data from control and ATF1/CREB1 dKO cells shows reads corresponding to *Crem* gene.

### ATF1/CREB1/CREM triple KO cells

We next produced ATF1/CREB1/CREM triple KO cells using CRISPR-Cas9 to create several CREM-deleted lines in one of the ATF1/CREB1 KO clones (**Figure 4**). As before, dDAVP produced large increases in AQP2 protein abundance in control cells, ATF1 single KO cells, and ATF1/CREB1 double KO cells. In contrast, the dDAVP response was markedly attenuated in the ATF1/CREB1/CREM triple KO cells (**Figure 4A and 4B**). We carried out RNA-seq analysis of dDAVP-treated ATF1/CREB1/CREM triple KO cells versus control cells with all three CREB-family transcription factors intact (**Figure 4C**). The curated data can be browsed, searched or downloaded at https://esbl.nhlbi.nih.gov/Databases/CREB-family/. A spreadsheet of all data values can be found at https://esbl.nhlbi.nih.gov/Databases/CREB-family/DataSets.html (Supplemental Spreadsheet 3). This analysis showed a striking decrease in *Aqp2* mRNA (upper left of **Figure 4C**), matching the effect on AQP2 protein. **Table 1** shows *Gene Ontology Biological Process* terms with over-representation of transcripts with altered abundance in ATF1/CREB1/CREM triple KO mpkCCD cells versus unmodified cells. Several of these terms refer to transport of ions and water, a central function of the CCD. Other terms (‘G protein-coupled receptor signaling pathway’ and ‘response to corticosteroid’) refer to signaling mechanisms in the renal collecting duct. Beyond this, several terms refer to processes that determine epithelial structure including ‘tube morphogenesis’, ‘cell projection morphogenesis’, ‘canonical Wnt signaling pathway’, and ‘negative regulation of cell population proliferation’. Such processes overlap known roles of vasopressin in native collecting ducts^37^.

**Figure 4.**
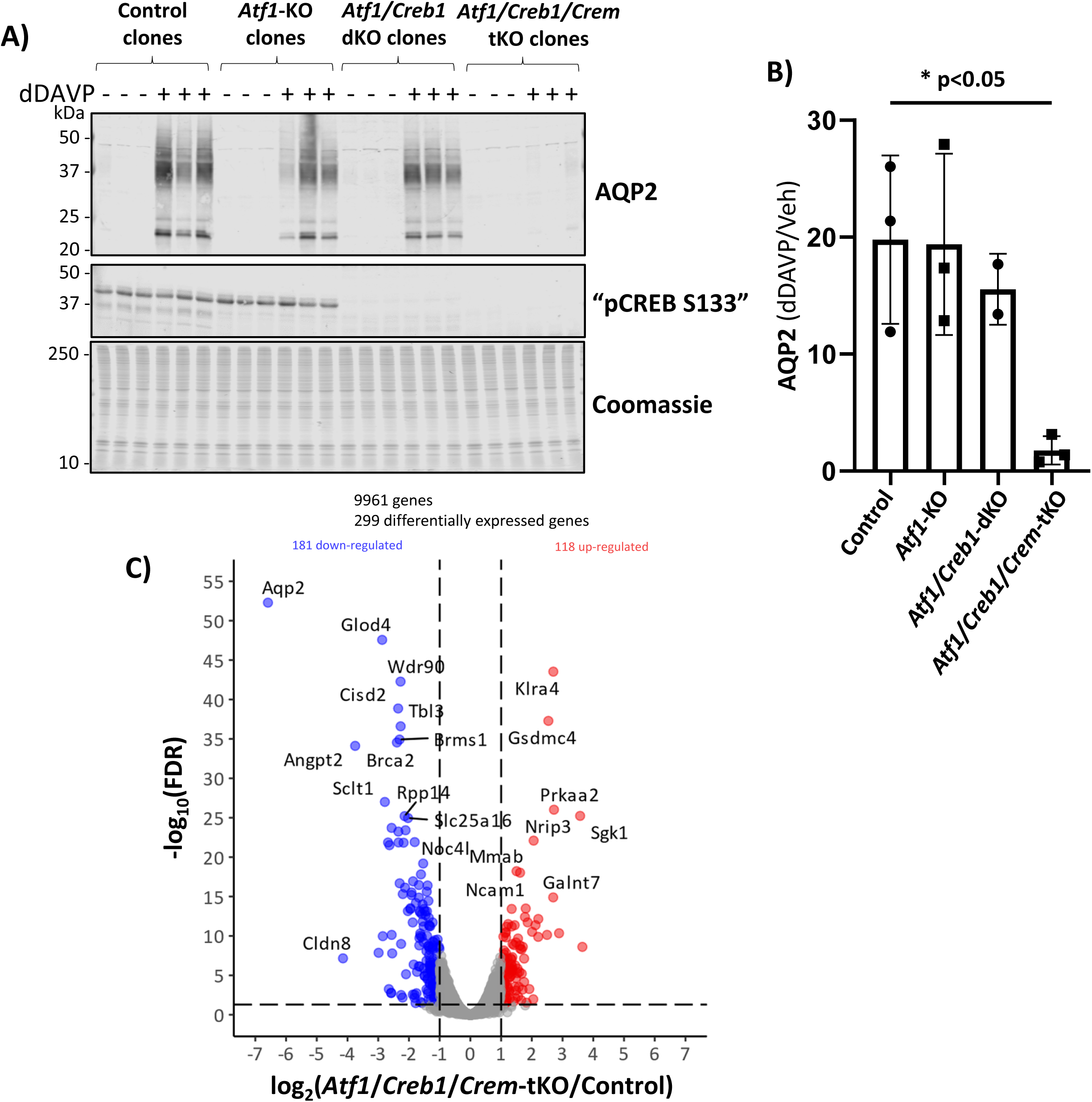
CRISPR-Cas9 deletion of ATF1, CREB1 and CREM1 in mouse mpkCCD cells. (**A**) Immunoblotting for AQP2, pCREB and staining with Coomassie blue in successive sub-panels. Observations were made in cells exposed to dDAVP (dDAVP +) of vehicle (dDAVP -) for 72 hours after confluence. The first subpanel show that AQP2 is upregulated by dDAVP. However, this response is decreased in ATF1/CREB1/CREM triple KO cells. Second subpanel shows ATF1/CREB1/CREM triple KO cells lack detectible signal with the pCREB antibody (which would recognize all three members of the family). (**B**) Densitometry for AQP2 (dDAVP +/dDAVP – ratio) in each group shows that the response is significantly lower only in the ATF1/CREB1/CREM triple KO cells. (**C**) RNA-seq volcano plot showing transcripts increased or decreased in ATF1/CREB1/CREM triple KO cells versus unmodified (control) cells according to threshold values: absolute value of log_2_(*Atf1/Creb1/Crem*-tKO/Control) greater than 1 and -log_10_(FDR-adjusted P value) greater than 1.301. All cells were grown in 0.1 nM dDAVP for 72 hours after confluence. Note that *Aqp2* mRNA is one of the top down-regulated genes.

**Table 1.**
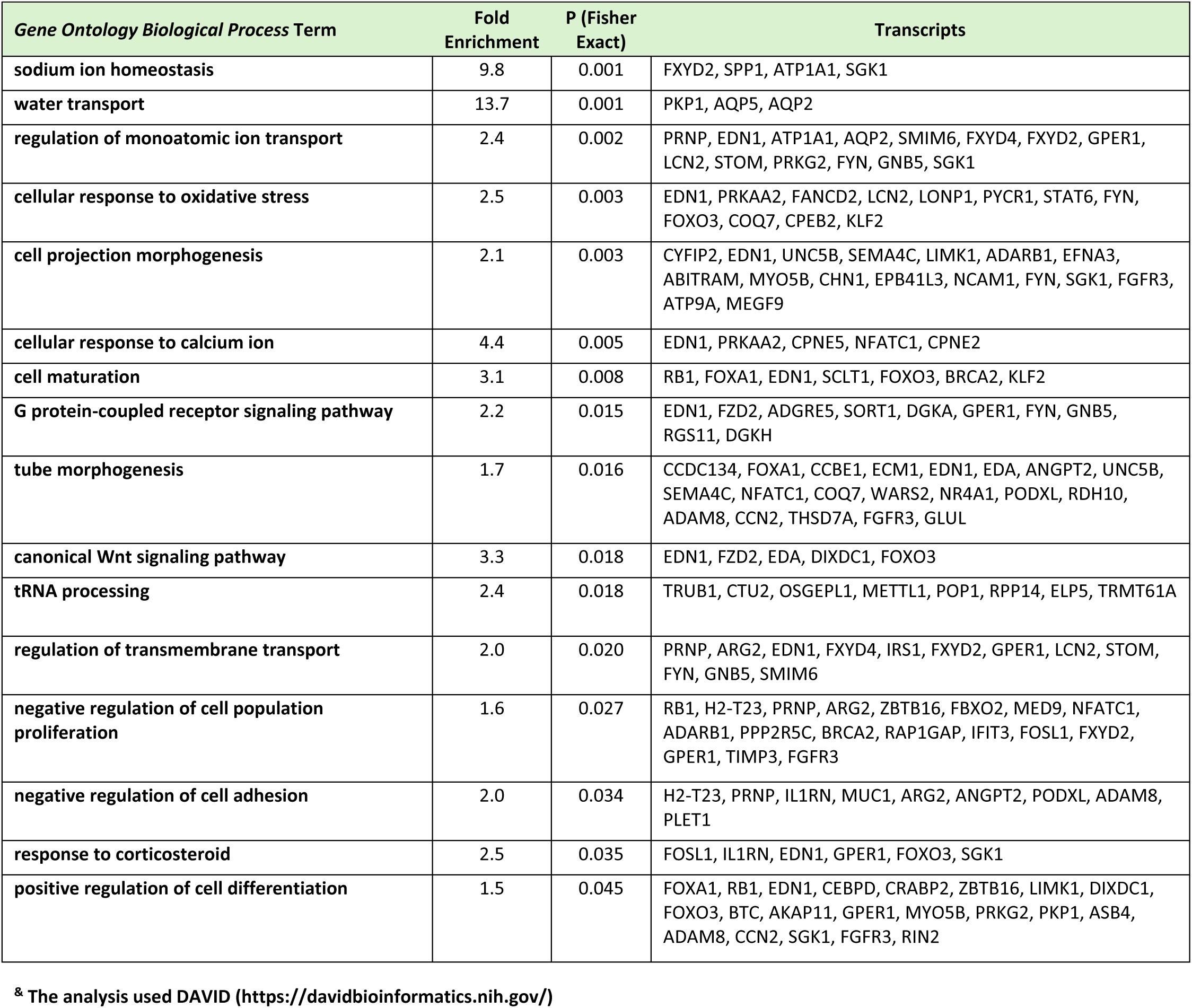
Gene Ontology Biological Process terms with over-representation of transcripts with significantly altered abundance in ATF1/CREB1/CREM triple KO mpkCCD cells vs. unmodified cells.^&^.

The volcano plot is similar to that seen in PKA double KO cells versus PKA intact cells^17^ in the sense that the *Aqp2* mRNA showed by far the most profound decrease among all mRNAs. In addition, prior studies quantifying changes in specific mRNAs in response to vasopressin show that *Aqp2* mRNA changes far more than any other transcript^16,36^. Several other transcripts show changes in the ATF1/CREB1/CREM triple KO cells as well as PKA-double KO cells^17^, and have opposite changes in response to vasopressin, suggesting that these transcripts are also regulated in the pathway triggered by the V2 receptor (**Table 2**). As seen in the ATF1/CREB1 double KO mpkCCD cells, the triple KO cells also showed an increase in the *Sgk1* transcript (**Figure 4C**) although prior studies did not demonstrate an effect of vasopressin to decrease *Sgk1* transcript^16^. Additionally, the RNA-seq analysis showed that ATF1/CREB1/CREM triple deletion resulted in significant abundance changes in multiple transcription factors (**Table 3**) and protein kinases (**Table 4**) that may play roles in the ATF1/CREB1/CREM -mediated regulation of *Aqp2* transcription by vasopressin.

**Table 2.**
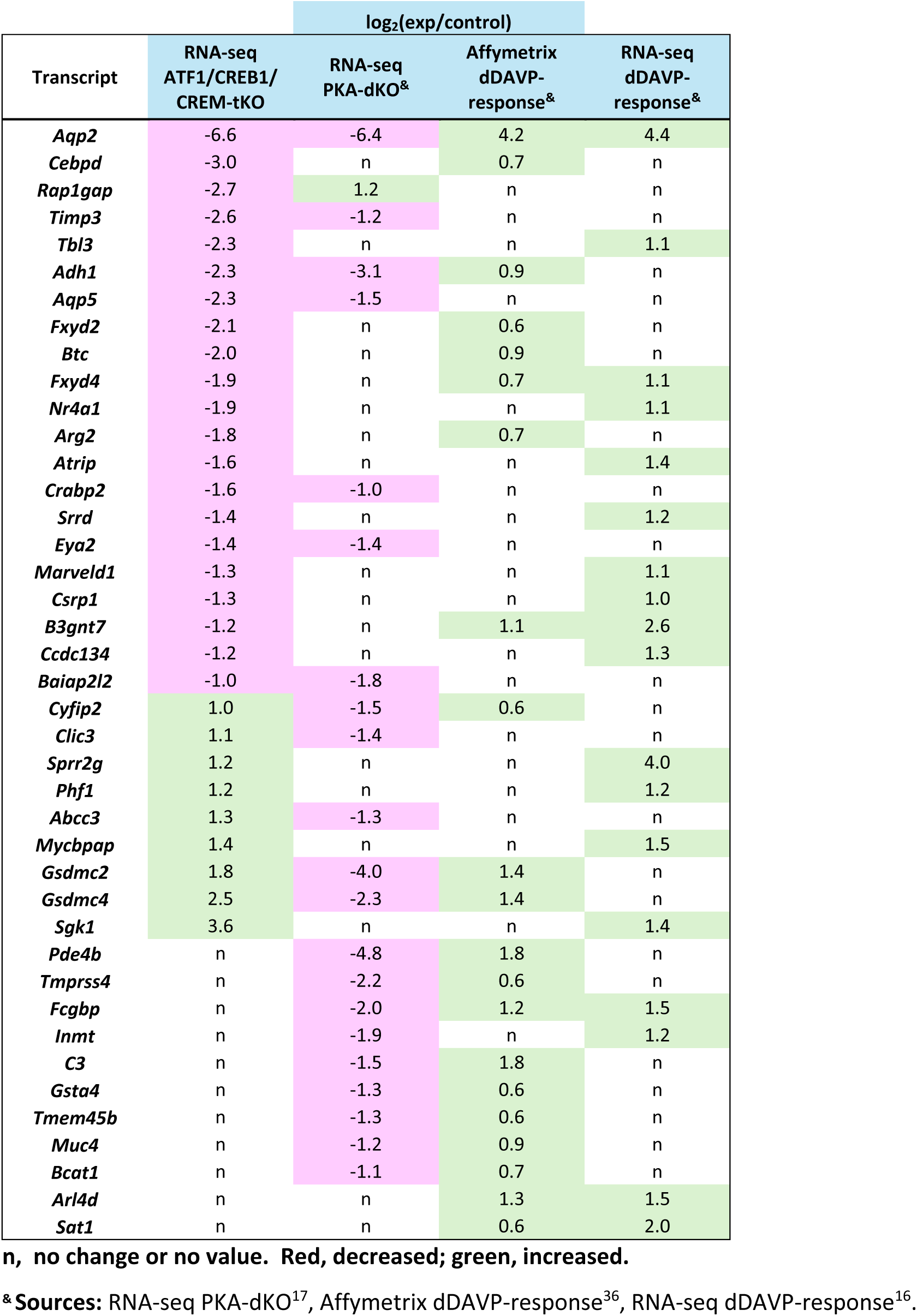
Comparison of transcriptomic findings from four studies.

**Table 3.** Transcription factors that undergo significant changes in ATF1/CREB1/CREM triple KO versus unmodified mpkCCD cells.

**A. Decreased in ATF1/CREB1/CREM triple KO cells**
| Gene symbol | Annotation | TPM ratio (Triple KO/ control) | log<sub>2</sub>(CPM) | P (FDR-Corrected) | TF Family<sup>&</sup> |
| --- | --- | --- | --- | --- | --- |
| *Camta2* | calmodulin binding transcription activator 2 | 0.45 | 4.42 | 1.10E-04 | CG-1 |
| *Cebpb* | CCAAT/enhancer binding protein (C/EBP), beta | 0.60 | 7.08 | 5.31E-05 | C/EBP |
| *Cebpd* | CCAAT/enhancer binding protein (C/EBP), delta | 0.13 | 4.52 | 1.37E-08 | C/EBP |
| *Etv3* | ets variant 3 | 0.57 | 5.49 | 4.97E-03 | ETS |
| *Fos* | FBJ osteosarcoma oncogene | 0.51 | 4.42 | 3.84E-05 | TF_bZIP |
| *Foxa1* | forkhead box A1 | 0.39 | 5.88 | 3.91E-12 | Fork head |
| *Hivep2* | human immunodeficiency virus type I enhancer binding protein 2 | 0.40 | 4.18 | 2.79E-08 | zf-C2H2 |
| *Jun* | jun proto-oncogene | 0.54 | 5.74 | 1.57E-03 | TF_bZIP |
| *Nr4a1* | nuclear receptor subfamily 4, group A, member 1 | 0.27 | 5.00 | 1.16E-17 | Nucl. receptor |
| *Plagl1* | pleiomorphic adenoma gene-like 1 | 0.22 | 4.98 | 4.80E-16 | zf-C2H2 |
| *Scx* | scleraxis | 0.40 | 5.23 | 1.31E-04 | bHLH |
| *Sox13* | SRY (sex determining region Y)-box 13 | 0.57 | 4.37 | 8.60E-04 | HMG |
| *Sox4* | SRY (sex determining region Y)-box 4 | 0.50 | 7.18 | 2.58E-07 | HMG |
| *Stat6* | signal transducer and activator of transcription 6 | 0.38 | 7.05 | 3.99E-17 | STAT |
| *Zfp219* | zinc finger protein 219 | 0.53 | 5.61 | 5.96E-06 | zf-C2H2 |
| *Zfp839* | zinc finger protein 839 | 0.51 | 4.65 | 7.25E-06 | zf-C2H2 |

**B. Increased in ATF1/CREB1/CREM triple KO cells**
| Gene symbol | Annotation | TPM ratio (Triple KO/ control) | log<sub>2</sub>(CPM) | P (FDR-Corrected) | TF Family<sup>&</sup> |
| --- | --- | --- | --- | --- | --- |
| *Foxn2* | forkhead box N2 | 1.69 | 4.24 | 5.98E-03 | Fork head |
| *Foxo3* | forkhead box O3 | 3.23 | 5.10 | 6.44E-04 | Fork head |
| *Id2* | inhibitor of DNA binding 2 | 1.87 | 7.70 | 3.91E-04 | bHLH |
| *Nfatc1* | nuclear factor of activated T cells, cytoplasmic, calcineurin dependent 1 | 2.20 | 4.81 | 8.24E-08 | RHD |
| *Tead4* | TEA domain family member 4 | 1.81 | 4.27 | 5.03E-04 | TEA |
| *Zbtb16* | zinc finger and BTB domain containing 16 | 2.41 | 4.31 | 7.05E-04 | ZBTB |
| *Zbtb8b* | zinc finger and BTB domain containing 8b | 1.92 | 5.07 | 9.32E-04 | ZBTB |
| *Zfp7* | zinc finger protein 7 | 1.71 | 7.36 | 2.61E-05 | zf-C2H2 |
| *Zfp763* | zinc finger protein 763 | 1.68 | 4.32 | 3.39E-03 | zf-C2H2 |

<sup>&</sup> C/EBP, CCAAT/Enhancer-Binding Proteins; ETS, E26 Transformation-Specific Factors; TF\_bZIP, Basic Leucine Zipper Transcription Factors; zf-C2H2, Zinc Finger C2H2 Transcription Factors; bHLH, Basic Helix-Loop-Helix Transcription Factors; CG-1; CG-1 Domain Transcription Factor Family; HMG, Uigh Mobility Group Transcription Factors; STAT, Signal Transducer and Activator of Transcription Proteins; RHD, Rel Homology Domain Transcription Factors; TEA, TEA Domain Transcription Factors; ZBTB, Zinc Finger and BTB Domain Transcription Factors.

**Table 4.** Protein kinases that undergo significant changes in ATF1/CREB1/CREM triple KO versus unmodified mpkCCD cells.

**A. Decreased in ATF1/CREB1/CREM triple KO cells**
| Gene symbol | Annotation | TPM ratio<br>(Triple KO/<br>control) | log<sub>2</sub>(CPM) | P (FDR-Corrected) | Kinase<br>Family<sup>&</sup> |
| --- | --- | --- | --- | --- | --- |
| *Camk1* | calcium/calmodulin-dependent protein kinase I | 0.56 | 5.77 | 1.49E-04 | CAMK |
| *Fgfr3* | fibroblast growth factor receptor 3 | 0.32 | 5.35 | 1.59E-09 | Tyr |
| *Mapkapk3* | mitogen-activated protein kinase-activated protein kinase 3 | 0.57 | 5.89 | 5.69E-04 | CAMK |
| *Nek1* | NIMA (never in mitosis gene a)-related expressed kinase 1 | 0.58 | 4.97 | 4.49E-04 | NEK |
| *Prkg2* | protein kinase, cGMP-dependent, type II | 0.38 | 4.42 | 2.50E-05 | AGC |
| *Snrk* | SNF related kinase | 0.47 | 4.80 | 2.02E-07 | CAMK |
| *Stk40* | serine/threonine kinase 40 | 0.56 | 4.83 | 3.18E-04 | CAMK |
| *Tnk1* | tyrosine kinase, non-receptor, 1 | 0.59 | 5.91 | 1.76E-04 | Tyr |

**B. Increased in ATF1/CREB1/CREM triple KO cells**
| Gene symbol | Annotation | TPM ratio<br>(Triple KO/<br>control) | log<sub>2</sub>(CPM) | P (FDR-Corrected) | Kinase<br>Family<sup>&</sup> |
| --- | --- | --- | --- | --- | --- |
| *Fyn* | Fyn proto-oncogene | 3.07 | 4.07 | 4.18E-09 | Tyr |
| *Hipk1* | homeodomain interacting protein kinase 1 | 1.76 | 8.32 | 4.89E-03 | CMGC |
| *Nrk* | Nik related kinase | 1.90 | 6.07 | 7.16E-04 | STE |
| *Plk3* | polo like kinase 3 | 1.85 | 5.31 | 5.48E-05 | Other |
| *Prkaa2* | protein kinase, AMP-activated, alpha 2 catalytic subunit | 6.59 | 4.76 | 9.53E-27 | CAMK |
| *Sgk1* | serum/glucocorticoid regulated kinase 1 | 11.88 | 9.66 | 5.83E-26 | AGC |
| *Stk38l* | serine/threonine kinase 38 like | 1.69 | 5.95 | 2.82E-04 | AGC |

<sup>&</sup>Tyr, Tyrosine Kinase Family; CAMK Calcium/Calmodulin-Dependent Protein Kinase-Related Family; NEK, NIMA-Related Kinase Family; AGC, Protein Kinase A/G/C Family; CMGC, CDK/MAPK/GSK3/CLK Kinase Family; STE, Sterile Kinase-Related Family

With this result we can conclude that either 1) CREM alone is responsible for the ability of dDAVP to increase *Aqp2* gene transcription; or 2) that any one of the three CREB-family transcription factors is sufficient to mediate the response to dDAVP and only when all three were deleted, did we see a diminution of the response. To discriminate between these two possibilities we carried out rescue experiments.

### Rescue experiments

Figure 5 reports the results of the rescue experiments in which the ATF1/CREB1/CEM triple KO cells underwent lentiviral mediated transduction with the three FLAG-epitope-tagged CREB-family transcription factors individually. All of the control clones showed a large increase in AQP2 protein abundance in response to dDAVP, whether or not they were transduced. Companion immunoblots using an anti-FLAG antibody demonstrated the success of the transduction (Figure 5A). In the triple KO cells, with no transduction, AQP2 was nearly undetectable with and without dDAVP exposure. In contrast, dDAVP elicited a large increase in AQP2 protein abundance following transduction of any of the three CREB-family transcription factors. Quantification of the response for all replicates (**Figure 5B**) confirms the finding that addition of any of the three transcription factors (ATF1 or CREB1 or CREM) is sufficient for a response to vasopressin.

**Figure 5.**
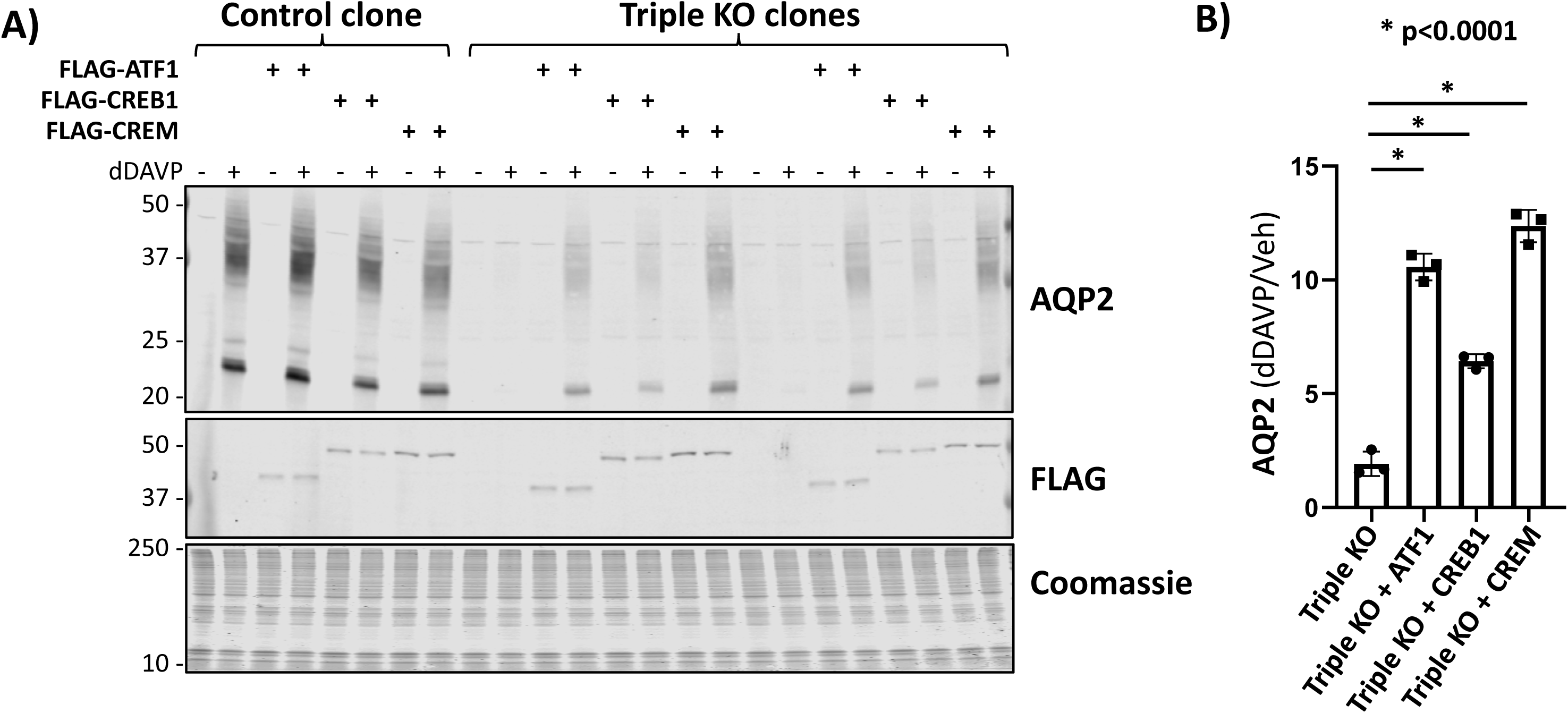
Rescue experiment in ATF1/CREB1/CREM-knockout cells. (**A**) Immunoblotting for AQP2, FLAG and staining with Coomassie blue in successive sub-panels. Observations were made in cells exposed to dDAVP (dDAVP +) of vehicle (dDAVP -) for 72 hours after confluence. The first subpanel show that AQP2 is upregulated by dDAVP in control cells with or without transduction. However, ATF1/CREB1/CREM triple KO cells show very low levels of AQP2 even in the presence of dDAVP. Re-expression of any one of the three CREB-related transcription factor promotes a rescue in the response of AQP2 abundance with vasopressin. Second subpanel shows the anti-FLAG antibody is able to recognize the corresponding tagged-transcription factors. (**B**) Densitometry for AQP2 (dDAVP +/dDAVP – ratio) in each group shows re-expression of any one of the three transcription factors causes a significantly higher response compared to un-transduced ATF1/CREB1/CREM triple KO cells.

## DISCUSSION

Vasopressin regulates water transport in the renal collecting duct in part through long term effects to accelerate transcription of the *Aqp2* gene, which codes for the aquaporin-2 water channel protein. Extensive studies in animal models of water balance disorders, including both water-retention syndromes and polyuric syndromes, indicated that these disorders are due largely to failure of the processes that regulate *Aqp2* gene transcription^8,38^. Hence progress in the diagnosis and treatment of water balance disorders depends on understanding the underlying mechanisms of *Aqp2* transcription and its regulation. An important sub-goal is to identify the transcription factors that regulate *Aqp2* transcription. The ‘conventional wisdom’ in the water balance field, repeated in multiple review articles, has been that the key transcription factor regulating *Aqp2* transcription is CREB, also known as CREB1 (reviewed in ***Introduction***). However, ChIP-seq studies identified no CREB binding sites within 390 kb of the *Aqp2* transcriptional start site, suggesting that CREB either plays no role in regulation of *Aqp2* transcription or only indirectly affects it^22^. That study raised doubt about the role of CREB. However, there are two other CREB-like transcription factors, ATF1 and CREM, that can be regulated by phosphorylation in a manner similar to CREB, which in principle could fulfill the proposed role of CREB.

The findings demonstrate that CREB1, ATF1 and CREM have redundant roles in regulation of *Aqp2* transcription and that only when all three were deleted in mpkCCD cells using CRISPR-Cas9 was the response to vasopressin to increase AQP2 expression attenuated. Rescue experiments in ATF1/CREB1/CREM triple KO cells showed that the response to vasopressin was restored when any of the three was expressed by lentivirus-mediated transduction, confirming the redundancy. The primary readout in these studies was immunoblotting for AQP2 protein although RNA-seq studies showed that *Aqp2* mRNA abundance was also altered in a manner parallel to AQP2 protein. Thus, the prior review articles proposing a role for CREB in regulation of *Aqp2* transcription were not totally wrong since expressing CREB alone was sufficient to restore the response to vasopressin in the ATF1/CREB1/CREM triple KO cells. In addition, the conclusion that there are no CREB1 binding sites near enough to regulate *Aqp2* transcription is likely to also be true for the other two CREB-like transcription factors because the bZIP DNA binding domains of the three are nearly identical (**Figure 6A, 6B and 6C**). Thus, we propose here (**Figure 6D**) that the three CREB-family transcription factors do not directly regulate *Aqp2* transcription, but rather work indirectly to regulate some other as-yet-unidentified transcription factors (possibly included in **Table 2**). We note also that the vasopressin response to increase AQP2 abundance was not totally abolished in the ATF1/CREB1/CREM triple KO cells (**Figure 4A and 4B**) indicating that there may be a small parallel response to vasopressin that is not dependent on any of the three CREB-family transcription factors.

**Figure 6.**
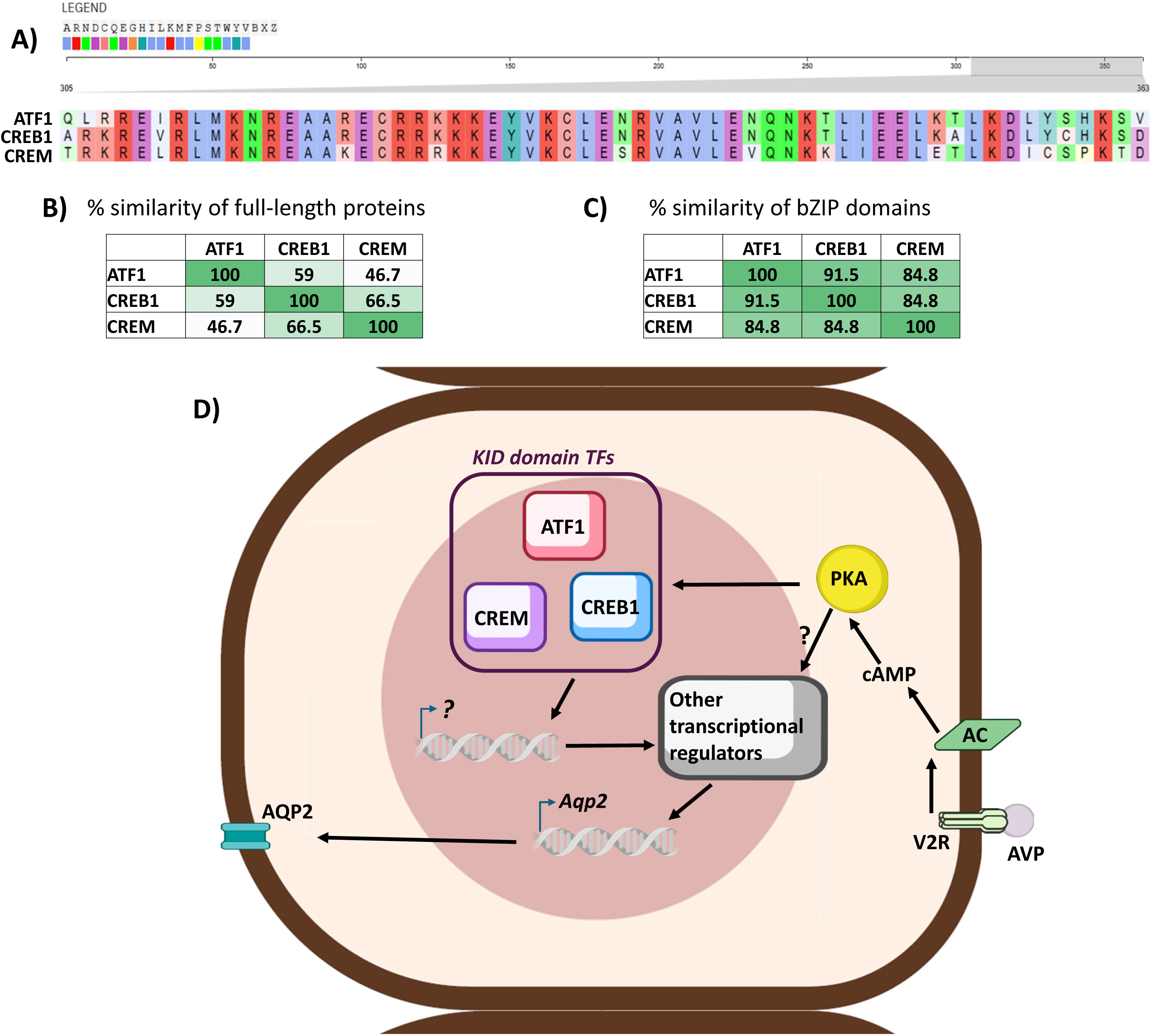
Proposed model of the role of CREB-family transcription factors in the regulation of AQP2 in response to vasopressin. (**A**) Sequence alignment of the bZIP domain of mouse ATF1, CREB1 and CREM. (**B**) Percentage similarity matrix considering the full-length sequence of the three transcription factors, showing a high degree of similarity among the three members. (**C**) Percentage similarity matrix only considering the bZIP domain of the three transcription factors, showing an even higher degree of similarity, suggesting similar functional roles of the DNA binding domain. (**D**) Cartoon model including the three members of the CREB-family transcription factors, characterized by the presence of KID domain. These transcription factors might regulate additional regulatory genes that ultimately lead to the increase in *Aqp2* gene transcription in the presence of vasopressin. Figure made using the NIH BIOART source (https://bioart.niaid.nih.gov/)

This study was carried out in a well-characterized cultured cell model of the cortical collecting duct, which recapitulates the characteristic short- and long-term responses to vasopressin to regulate AQP2 (**Introduction**)^33^. The strength of this model compared with doing similar studies in mice is the ease of genetic manipulation. The complexity involved in doing multiple inducible, collecting-duct-targeted gene deletions in mice is daunting. Thus, the mpkCCD model fulfilled the role of facilitating cellular level studies that would be virtually impossible in intact mice. Future studies will be needed to further identify binding sites for ATF1, CREB1, and CREM along the genome, recognizing that the profiles of ATF1 and CREM binding are likely to be similar, if not identical, to that of CREB1^22^ because of the marked similarity of the respective bZIP DNA-binding domains in the three transcription factors (**Figure 6A, 6B and 6C)**. In addition, if the effects of the CREB-family transcription factors on *Aqp2* gene transcription are indirect as proposed in **Figure 6D**, then future studies (e.g. CRISPR screening) will be needed to identify the transcription factors that directly mediate the regulation.

## ACKNOWLEDGMENTS

This research was supported in part by the Intramural Research Program of the NIH and NHLBI. Cell sorting was performed in the NHLBI Flow Cytometry Core Facility (Dr. Pradeep Dagur, Director). Next-generation sequencing was done in the NHLBI DNA Sequencing Core Facility (Yuesheng Li, Director).

## DATA SHARING

Sequencing data have been deposited in the GEO: https://www.ncbi.nlm.nih.gov/geo/query/acc.cgi?acc=GSE291084

(secure token is available upon request).

Summaries of curated RNA-seq data are available in supplemental spreadsheets available at: https://esbl.nhlbi.nih.gov/Databases/CREB-family/DataSets.html.

## AUTHOR CONTRIBUTIONS

Conception and Designed research: A.R.M.-D.-O., M.A.K.; conducted experiments: A.R.M.-D.-O., L.C., S.- M.O., E.P. S.K.; analyzed data: A.R.M.-D.-O., L.C., S.-M.O., V.R., C.-R.Y., C.-L.C., M.A.K.; interpreted results of experiments: A.R.M.-D.-O., L.C., V.R., C.-R.Y., C.-L.C., M.A.K.; prepared figures: A.R.M.-D.-O., M.A.K.; drafted manuscript: A.R.M.-D.-O., M.A.K.; edited and revised manuscript: all authors; approved final version of manuscript: all authors.

## SUPPLEMENTAL MATERIAL

**Supplemental Spreadsheet 1. RNA-seq comparison of transcript abundances in ATF1 single knockout mpkCCD cells and control cells without the deletion.**

**Supplemental Spreadsheet 2. RNA-seq comparison of transcript abundances in ATF1/CREB1 double knockout mpkCCD cells and control cells without the deletions.**

**Supplemental Spreadsheet 3. RNA-seq comparison of transcript abundances in ATF1/CREB1/CREM triple knockout mpkCCD cells and control cells without the deletions.**

**Available for download at: https://esbl.nhlbi.nih.gov/Databases/CREB-family/DataSets.html**

## REFERENCES

1. Knepper MA, Kwon TH, Nielsen S. Molecular physiology of water balance. N Engl J Med. Apr 2 2015;372(14):1349–1358. doi:10.1056/NEJMra1404726

2. D’Acierno M, Fenton RA, Hoorn EJ. The biology of water homeostasis. Nephrol Dial Transplant. Oct 21 2024;doi:10.1093/ndt/gfae235

3. Olesen ETB, Fenton RA. Aquaporin 2 regulation: implications for water balance and polycystic kidney diseases. Nat Rev Nephrol. Nov 2021;17(11):765–781. doi:10.1038/s41581-021-00447-x

4. Cheung PW, Bouley R, Brown D. Targeting the Trafficking of Kidney Water Channels for Therapeutic Benefit. Annu Rev Pharmacol Toxicol. Jan 6 2020;60:175–194. doi:10.1146/annurev-pharmtox-010919-023654

5. Pearce D, Soundararajan R, Trimpert C, Kashlan OB, Deen PM, Kohan DE. Collecting duct principal cell transport processes and their regulation. Clin J Am Soc Nephrol. Jan 7 2015;10(1):135–146. doi:10.2215/CJN.05760513

6. Kortenoeven ML, Pedersen NB, Rosenbaek LL, Fenton RA. Vasopressin regulation of sodium transport in the distal nephron and collecting duct. Am J Physiol Renal Physiol. Aug 15 2015;309(4):F280–299. doi:10.1152/ajprenal.00093.2015

7. Ecelbarger CA, Kim GH, Terris J, et al. Vasopressin-mediated regulation of epithelial sodium channel abundance in rat kidney. Am J Physiol Renal Physiol. Jul 2000;279(1):F46–53. doi:10.1152/ajprenal.2000.279.1.F46

8. Nielsen S, Frokiaer J, Marples D, Kwon TH, Agre P, Knepper MA. Aquaporins in the kidney: from molecules to medicine. Physiol Rev. Jan 2002;82(1):205–244. doi:10.1152/physrev.00024.2001

9. Verbalis JG. Disorders of body water homeostasis. Best Pract Res Clin Endocrinol Metab. Dec 2003;17(4):471–503. doi:10.1016/s1521-690x(03)00049-6

10. Valenti G, Tamma G. The vasopressin-aquaporin-2 pathway syndromes. Handb Clin Neurol. 2021;181:249–259. doi:10.1016/B978-0-12-820683-6.00018-X

11. Verbalis JG. Disorders of water metabolism: diabetes insipidus and the syndrome of inappropriate antidiuretic hormone secretion. Handb Clin Neurol. 2014;124:37–52. doi:10.1016/B978-0-444-59602-4.00003-4

12. Rosner MH, Rondon-Berrios H, Sterns RH. Syndrome of Inappropriate Antidiuresis. J Am Soc Nephrol. Dec 2 2024;doi:10.1681/ASN.0000000588

13. Nielsen S, Chou CL, Marples D, Christensen EI, Kishore BK, Knepper MA. Vasopressin increases water permeability of kidney collecting duct by inducing translocation of aquaporin-CD water channels to plasma membrane. Proc Natl Acad Sci U S A. Feb 14 1995;92(4):1013–1017. doi:10.1073/pnas.92.4.1013

14. DiGiovanni SR, Nielsen S, Christensen EI, Knepper MA. Regulation of collecting duct water channel expression by vasopressin in Brattleboro rat. Proc Natl Acad Sci U S A. Sep 13 1994;91(19):8984–8988. doi:10.1073/pnas.91.19.8984

15. Matsumura Y, Uchida S, Rai T, Sasaki S, Marumo F. Transcriptional regulation of aquaporin-2 water channel gene by cAMP. J Am Soc Nephrol. Jun 1997;8(6):861–867. doi:10.1681/ASN.V86861

16. Sandoval PC, Claxton JS, Lee JW, Saeed F, Hoffert JD, Knepper MA. Systems-level analysis reveals selective regulation of Aqp2 gene expression by vasopressin. Sci Rep. Oct 11 2016;6:34863. doi:10.1038/srep34863

17. Isobe K, Jung HJ, Yang CR, et al. Systems-level identification of PKA-dependent signaling in epithelial cells. Proc Natl Acad Sci U S A. Oct 17 2017;114(42):E8875–E8884. doi:10.1073/pnas.1709123114

18. Kortenoeven ML, Fenton RA. Renal aquaporins and water balance disorders. Biochim Biophys Acta. May 2014;1840(5):1533–1549. doi:10.1016/j.bbagen.2013.12.002

19. Bockenhauer D, Bichet DG. Pathophysiology, diagnosis and management of nephrogenic diabetes insipidus. Nat Rev Nephrol. Oct 2015;11(10):576–588. doi:10.1038/nrneph.2015.89

20. Hozawa S, Holtzman EJ, Ausiello DA. cAMP motifs regulating transcription in the aquaporin 2 gene. Am J Physiol. Jun 1996;270(6 Pt 1):C1695-1702. doi:10.1152/ajpcell.1996.270.6.C1695

21. Yasui M, Zelenin SM, Celsi G, Aperia A. Adenylate cyclase-coupled vasopressin receptor activates AQP2 promoter via a dual effect on CRE and AP1 elements. Am J Physiol. Apr 1997;272(4 Pt 2):F443-450. doi:10.1152/ajprenal.1997.272.4.F443

22. Jung HJ, Raghuram V, Lee JW, Knepper MA. Genome-Wide Mapping of DNA Accessibility and Binding Sites for CREB and C/EBPbeta in Vasopressin-Sensitive Collecting Duct Cells. J Am Soc Nephrol. May 2018;29(5):1490–1500. doi:10.1681/ASN.2017050545

23. Mayr B, Montminy M. Transcriptional regulation by the phosphorylation-dependent factor CREB. Nat Rev Mol Cell Biol. Aug 2001;2(8):599–609. doi:10.1038/35085068

24. Johannessen M, Moens U. Multisite phosphorylation of the cAMP response element-binding protein (CREB) by a diversity of protein kinases. Front Biosci. Jan 1 2007;12:1814–1832. doi:10.2741/2190

25. Chrivia JC, Kwok RP, Lamb N, Hagiwara M, Montminy MR, Goodman RH. Phosphorylated CREB binds specifically to the nuclear protein CBP. Nature. Oct 28 1993;365(6449):855-859. doi:10.1038/365855a0

26. Molina CA, Foulkes NS, Lalli E, Sassone-Corsi P. Inducibility and negative autoregulation of CREM: an alternative promoter directs the expression of ICER, an early response repressor. Cell. Dec 3 1993;75(5):875–886. doi:10.1016/0092-8674(93)90532-u

27. Kikuchi H, Jung HJ, Raghuram V, et al. Bayesian identification of candidate transcription factors for the regulation of Aqp2 gene expression. Am J Physiol Renal Physiol. Sep 1 2021;321(3):F389–F401. doi:10.1152/ajprenal.00204.2021

28. Isobe K, Raghuram V, Krishnan L, Chou CL, Yang CR, Knepper MA. CRISPR-Cas9/phosphoproteomics identifies multiple noncanonical targets of myosin light chain kinase. Am J Physiol Renal Physiol. Mar 1 2020;318(3):F600–F616. doi:10.1152/ajprenal.00431.2019

29. Pisitkun T, Hoffert JD, Saeed F, Knepper MA. NHLBI-AbDesigner: an online tool for design of peptide-directed antibodies. Am J Physiol Cell Physiol. Jan 1 2012;302(1):C154–164. doi:10.1152/ajpcell.00325.2011

30. Hoffert JD, Fenton RA, Moeller HB, et al. Vasopressin-stimulated increase in phosphorylation at Ser269 potentiates plasma membrane retention of aquaporin-2. J Biol Chem. Sep 5 2008;283(36):24617–24627. doi:10.1074/jbc.M803074200

31. Chen L, Chou CL, Knepper MA. A Comprehensive Map of mRNAs and Their Isoforms across All 14 Renal Tubule Segments of Mouse. J Am Soc Nephrol. Apr 2021;32(4):897–912. doi:10.1681/ASN.2020101406

32. Chen L, Lee JW, Chou CL, et al. Transcriptomes of major renal collecting duct cell types in mouse identified by single-cell RNA-seq. Proc Natl Acad Sci U S A. Nov 14 2017;114(46):E9989–E9998. doi:10.1073/pnas.1710964114

33. Yu MJ, Miller RL, Uawithya P, et al. Systems-level analysis of cell-specific AQP2 gene expression in renal collecting duct. Proc Natl Acad Sci U S A. Feb 17 2009;106(7):2441–2446. doi:10.1073/pnas.0813002106

34. Sheng M, Thompson MA, Greenberg ME. CREB: a Ca(2+)-regulated transcription factor phosphorylated by calmodulin-dependent kinases. Science. Jun 7 1991;252(5011):1427-1430. doi:10.1126/science.1646483

35. Du K, Montminy M. CREB is a regulatory target for the protein kinase Akt/PKB. J Biol Chem. Dec 4 1998;273(49):32377–32379. doi:10.1074/jbc.273.49.32377

36. Khositseth S, Pisitkun T, Slentz DH, et al. Quantitative protein and mRNA profiling shows selective post-transcriptional control of protein expression by vasopressin in kidney cells. Mol Cell Proteomics. Jan 2011;10(1):M110 004036. doi:10.1074/mcp.M110.004036

37. Salhadar K, Matthews A, Raghuram V, et al. Phosphoproteomic Identification of Vasopressin/cAMP/Protein Kinase A-Dependent Signaling in Kidney. Mol Pharmacol. May 2021;99(5):358–369. doi:10.1124/mol.120.119602

38. Mak A, Sung CC, Pisitkun T, Khositseth S, Knepper MA. ’Aquaporin-omics’: mechanisms of aquaporin-2 loss in polyuric disorders. J Physiol. Jul 2024;602(13):3191–3206. doi:10.1113/JP284634

